# Parkin drives pS65-Ub turnover independently of canonical autophagy in *Drosophila*

**DOI:** 10.1101/2021.06.10.447841

**Authors:** Joanne L. Usher, Juliette J. Lee, Alvaro Sanchez-Martinez, Alexander J. Whitworth

## Abstract

Parkinson’s disease-related proteins, PINK1 and Parkin, act in a common pathway to maintain mitochondrial quality control. While the PINK1-Parkin pathway can promote autophagic mitochondrial turnover (mitophagy) in cell culture, recent studies have questioned whether they contribute to mitophagy *in vivo*, and alternative PINK1- and Parkin-dependent mitochondrial quality control pathways have been proposed. To determine the mechanisms by which the Pink1-Parkin pathway operates *in vivo*, we developed methods to detect Ser65-phosphorylated ubiquitin (pS65-Ub) in *Drosophila*. Exposure to the oxidant paraquat led to robust, Pink1-dependent pS65-Ub production. Surprisingly, *parkin*-null flies displayed strikingly elevated basal levels of pS65-Ub, suggestive of disrupted flux through the Pink1-parkin pathway. Depletion of the core autophagy proteins Atg1, Atg5 and Atg8a did not cause pS65-Ub accumulation to the same extent as loss of parkin, and overexpression of parkin was able to reduce both basal and paraquat-induced pS65-Ub levels in an *Atg5*-null background. Taken together, these results suggest that the Pink1-parkin pathway is able to promote mitochondrial turnover independently of canonical autophagy *in vivo*.

## Introduction

Parkinson’s disease (PD) is the second most common neurodegenerative disease, with the global burden of disease having more than doubled between 1990 and 2016 (GBD 2016 Parkinson’s Disease Collaborators et al., 2018). Autosomal recessive mutations in the genes encoding the mitochondria-targeted kinase PINK1 and the E3 ubiquitin (Ub) ligase Parkin are associated with parkinsonism (Kitada et al., 1998; Valente et al., 2004). Loss of either homolog in *Drosophila* (Pink1 and parkin, respectively) results in strikingly similar phenotypes of severe mitochondrial dysfunction and degeneration of the indirect flight muscles, as well as the degeneration of a subset of dopaminergic neurons, thus mimicking a key hallmark of PD (Greene et al., 2003; Whitworth et al., 2005; Clark et al., 2006; Park et al., 2006). Genetic interaction studies subsequently placed *Pink1* and *parkin* in a common pathway, with *parkin* downstream of *Pink1* (Clark et al., 2006).

The best-characterised PINK1 substrates are Ub and Parkin, each phosphorylated at their respective Ser65 residues (Kane et al., 2014; Kazlauskaite et al., 2014; Koyano et al., 2014; Kondapalli et al., 2012; Shiba-Fukushima et al., 2012). PINK1 is partially imported into healthy mitochondria via its N-terminal mitochondrial targeting sequence where it is cleaved and degraded in the cytosol by the N-end rule pathway (Yamano and Youle, 2013). PINK1 is activated upon stalling on the outer mitochondrial membrane (OMM), where it phosphorylates Ub (pS65-Ub) that is conjugated at low abundance to OMM proteins (Okatsu et al., 2015a). Parkin exists in the cytosol in an autoinhibited state and is recruited to mitochondria by binding pS65-Ub (Okatsu et al., 2015b). pS65-Ub binding partially displaces Parkin’s Ubl domain, which allows it to be phosphorylated by PINK1 (Gladkova et al., 2018). This second phosphorylation event results in a dramatic domain rearrangement that relieves Parkin’s autoinhibitory contacts and allows it to ubiquitinate proteins in close proximity (Gladkova et al., 2018; Sauvé et al., 2018). The Ub provided by Parkin allows further phosphorylation by PINK1, which in turn promotes further Parkin recruitment, thus constituting a feed-forward mechanism of mitochondrial pS65-ubiquitination that is dependent on both PINK1 and Parkin (Ordureau et al., 2014). Both the structure of active Parkin and cell-based studies suggest that Parkin has low substrate selectivity (Gladkova et al., 2018; Koyano et al., 2019), and it has been found to predominantly produce K6, K11, K48 and K63 chains *in vitro* (Ordureau et al., 2014).

Much of our understanding of the function of PINK1 and Parkin utilised chemical depolarisation of mitochondria in cultured cells in conjunction with Parkin overexpression (Narendra et al., 2008; Vives-Bauza et al., 2010). These experiments established a paradigm in studying the PINK1-parkin pathway; upon depolarisation, PINK1- and Parkin-mediated ubiquitination of OMM proteins leads to the recruitment of the Ub-binding mitophagy receptors OPTN and NDP52 (Lazarou et al., 2015), which in turn promote autophagosome initiation (Yamano et al., 2020; Boyle et al., 2019), ultimately leading to degradation of the damaged mitochondria via the autophagy system. However, studies in animal models have provided mixed results as to the contribution of PINK1 and Parkin to mitophagy as measured by pH-sensitive fluorescent reporter constructs (Lee et al., 2018; Cornelissen et al., 2018; McWilliams et al., 2018; Kim et al., 2019; Liu et al., 2021). It has also been shown in cell culture models that treatment with Antimycin A or expression of an aggregation-prone matrix protein, ∆OTC, neither of which cause mitochondrial depolarisation, led to the production of mitochondria-derived vesicles (MDVs) in a PINK1- and Parkin-dependent manner (McLelland et al., 2014; Burman et al., 2017). Other studies have focused on the role of PINK1 and Parkin in mitochondrial biogenesis, protein import, and in the regulation of the fission and fusion machinery (Stevens et al., 2015; Jacoupy et al., 2019; Poole et al., 2008). However, many questions remain about the mechanisms of PINK1-Parkin-mediated mitochondrial quality control *in vivo*.

We sought to determine the physiological mechanisms of the PINK1-parkin pathway by monitoring pS65-Ub levels as a direct measure of PINK1 activity. We developed complementary mass spectrometry, immunoblotting and immunostaining methods to detect pS65-Ub using *Drosophila* as a model system. We confirm that pS65-Ub production is Pink1-dependent and can therefore be utilised to follow activation of the Pink1-parkin pathway, and the downstream mechanisms of mitochondrial turnover, *in vivo*. We identify exposure to the oxidant and parkinsonian toxin paraquat as a potent activator of the Pink1-parkin pathway, and establish this approach as a new paradigm to study the Pink1-parkin pathway *in vivo*.

## Results

### Development of methods to detect pS65-Ub in vivo

To understand the role of the Pink1-parkin pathway in maintaining mitochondrial quality control *in vivo*, we developed methods to detect pS65-Ub at low abundances by mass spectrometry. Using a sample preparation pipeline based on the recently described Ub-Clipping method (Swatek et al., 2019), we determined the absolute abundance of total Ub and pS65-Ub in mitochondrial extracts from young (2-3 days) and aged (50 to 60 days) flies from a wild-type background (*w*^1118^). In young flies, Ub was present on mitochondria, but pS65-Ub was not reliably detected with this method (Figure 1A). In contrast, aged flies displayed elevated total mitochondrial Ub, and we were able to robustly detect pS65-Ub (Figure 1A). To gain a clearer insight into the basal levels of pS65-Ub in young wild type animals, we adjusted our approach in order to optimise pS65-Ub detection (see Methods). Using this method we were indeed able to detect pS65-Ub in mitochondrial fractions from young flies (Figure 1B), thus confirming that pS65-Ub is present in young flies at very low abundance.

**Figure 1:**
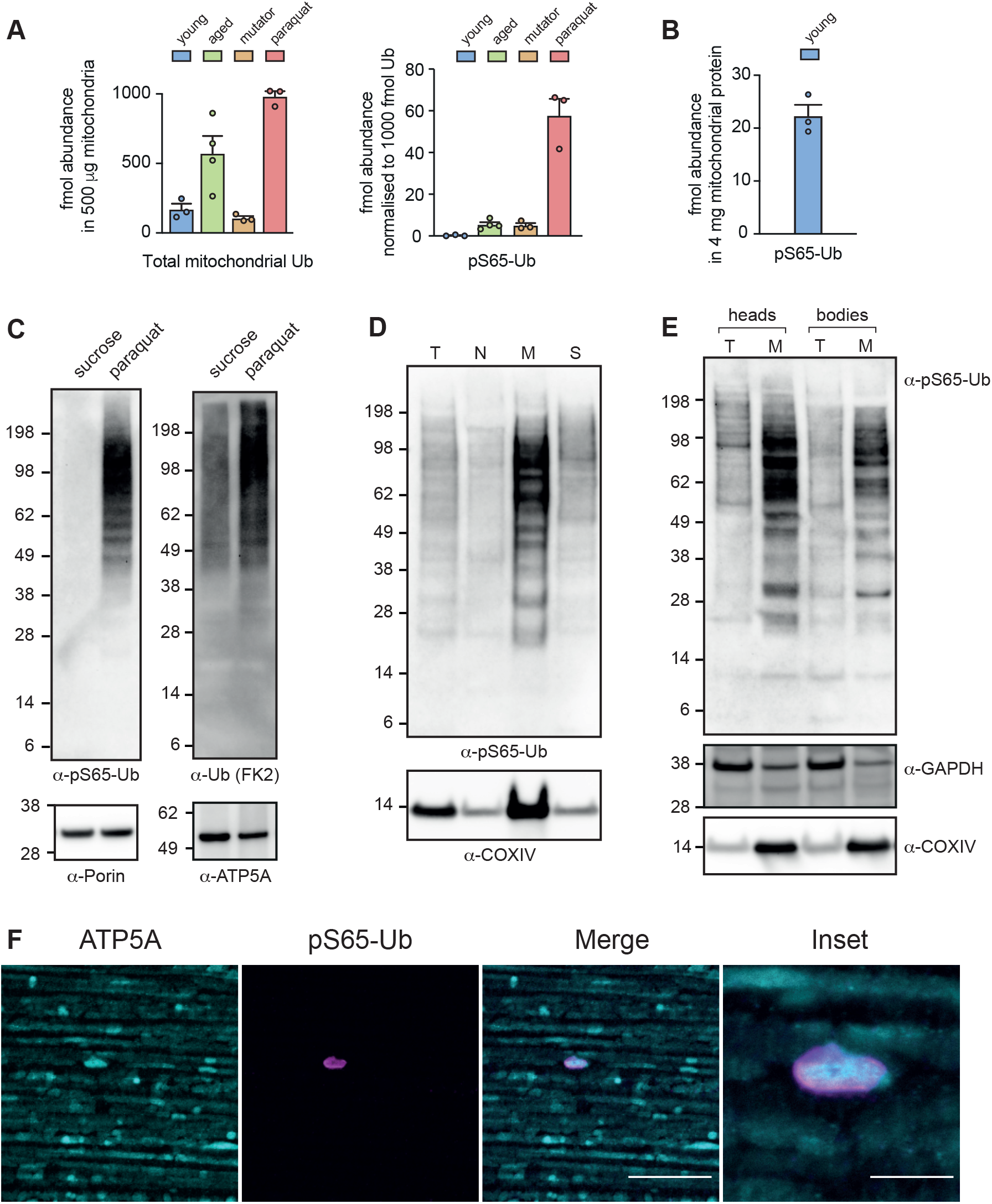
Detection of pS65-Ub *in vivo*. (A) Total Ub (left) and normalised pS65-Ub abundance (right) in 500 μg Ub-Clippase-treated, TUBE-enriched mitochondrial fractions from young (2-3 days) and aged (50-60 days) wild-type flies, an mtDNA mutator model (*daG4>UAS-mito*-*APOBEC1*), and wild-type flies that had been exposed to paraquat (5 mM) for 3 days. (B) Absolute abundance of pS65-Ub in 4 mg mitochondria from young (2-3 days) wild-type flies following sodium carbonate extraction, Ub-Clippase treatment and phospho-peptide enrichment. Charts show mean +/− SEM, n = 3-4 independent biological replicates as shown. (C) pS65-Ub (left) and total Ub (right) immunoblots of mitochondria-enriched fractions from wild-type flies treated 3 days with either paraquat or vehicle (sucrose). (D) pS65-Ub immunoblot following subcellular fractionation of flies treated with paraquat for 3 days. T, total lysate; N, nuclear-enriched fraction; M, mitochondria-enriched fraction; S, post-mitochondrial supernatant. (E) pS65-Ub immunoblot following mechanical separation of fly heads from bodies (thoraces and abdomens). (F) Representative image of flight muscles from aged (50-day-old) wild-type animals immunostained with the indicated antibodies. Scale bars = 20 μm (inset, 5 μm).

We next sought to identify *in vivo* stimuli of the Pink1-parkin pathway. Mitochondrial stress induced by a mtDNA mutator model (*daG4*>*mito*-*APOBEC1*) (Andreazza et al., 2019) was sufficient to induce pS65-Ub in young flies (Figure 1A). We then tested paraquat, an oxidant that has been epidemiologically linked to PD (Tanner et al., 2011). In striking contrast to the ageing and mutator models, exposure to paraquat led to a robust increase in both total and pS65-Ub in mitochondrial fractions (Figure 1A). In paraquat-treated flies, pS65-Ub comprised approximately 6% of the total mitochondrial Ub, substantially less than the 10-30% Ub phosphorylation that has been observed using similar methods in depolarised cells (Ordureau et al., 2014; Swatek et al., 2019).

As an orthogonal validation of our mass spectrometry results, we evaluated immunodetection methods using an antibody recently characterised to specifically detect pS65-Ub at the femtomolar to picomolar range (Watzlawik et al., 2020). Immunoblotting confirmed the robust induction of pS65-Ub and total mitochondrial Ub in response to paraquat, while pS65-Ub was not detected in response to amino acid starvation from a sucrose-only diet (Figure 1C). The paraquat-induced pS65-Ub co-enriched with mitochondria in sub-cellular fractions (Figure 1D). Interestingly, pS65-Ub levels appeared to be greater in heads compared with bodies (thorax and abdomens) (Figure 1E). We also determined that removal of paraquat led to a reduction in pS65-Ub levels, presumably due to mitochondrial turnover (Supplementary Figure 1A). Immunofluorescence microscopy of the flight muscles of aged flies revealed low but consistent detection of mitochondria (ATP5A-positive) that were enveloped in pS65-Ub (Figure 1F), while pS65-Ub-positive structures were rarely observed in young flies (Supplementary Figure 2A). These results suggest that the Pink1-parkin pathway is basally active in *Drosophila*, but that pS65-Ub levels are likely kept very low in young animals due to efficient turnover.

### pS65-Ub production in response to paraquat requires Pink1 but not parkin

We next sought to confirm the requirements for Pink1 and parkin in the ubiquitination of mitochondria under basal and paraquat-induced conditions. To this end, we determined the abundance of total Ub and pS65-Ub in mitochondrial extracts from wild-type, *Pink1*^−^ (*Pink1*^B9^), and *park*^−/−^ (*park*^25^) mutant flies by mass spectrometry. Under basal conditions, loss of Pink1 resulted in elevated total Ub levels that did not further increase upon exposure to paraquat (Figure 2A). Importantly, pS65-Ub levels did not increase above background even upon exposure to paraquat in *Pink1*^−^ flies (Figure 2B, Supplementary Figure 1B), confirming the conserved and essential role of Pink1 in the phosphorylation of Ub at Ser65 in *Drosophila*. *park*^−/−^ flies displayed modestly elevated total mitochondrial Ub that did not significantly increase in response to paraquat (Figure 2A). In contrast, the increase in pS65-Ub levels observed upon exposure to paraquat was, surprisingly, largely unaffected by loss of parkin (Figure 2B).

**Figure 2:**
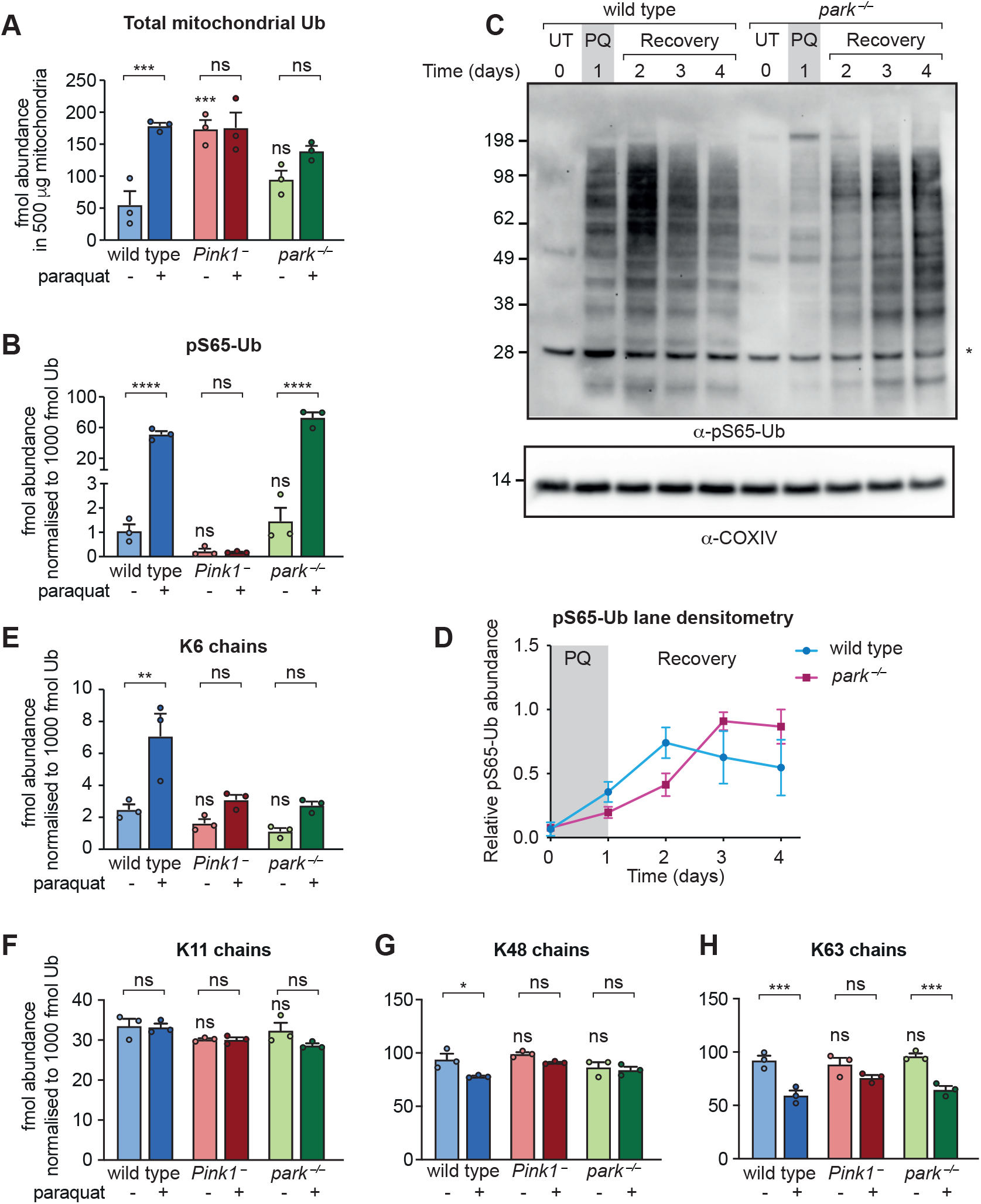
Analysis of the paraquat-induced mitochondrial ubiquitome from *Pink1*^−^ and *park*^−/−^ flies. (A) total Ub and (B) normalised pS65-Ub abundance in 500 μg TUBE-enriched, Ub-Clippase-treated mitochondrial fractions from wild-type, *Pink1*^−^ and *park*^−/−^ flies, either untreated young flies (2-3 days) or exposed to paraquat for 3 days. (C) pS65-Ub immunoblot of mitochondria-enriched fractions following the paraquat pulse-chase assay in wild-type and *park*^−/−^ flies. UT, untreated; PQ, 1-day paraquat treatment; Recovery, flies removed from paraquat and returned to normal food. * = non-specific band. (D) pS65-Ub lane densitometry analysis of n = 3 independent replicates from (C), expressed relative to the most intense lane signal in each blot. Charts show mean +/− SEM. (E-H) Relative abundance of (E) K6 chains, (F) K11 chains, (G) K48 chains, and (H) K63 chains in mitochondrial fractions treated as in A, normalised to the total mitochondrial Ub in each sample. Charts show mean +/− SEM from n = 3 independent biological replicates. Statistical analysis used one-way ANOVA with Šidák’s correction for multiple comparisons. * *P* < 0.05; ** *P* < 0.01; *** *P* < 0.001; **** *P* < 0.0001, ns = non-significant. The full list of multiplicity-adjusted *P* values is presented in Supplementary Table 1.

As an intermediate product in the Pink1-parkin pathway, pS65-Ub levels will be affected by the kinetics of both its production and downstream turnover, both of which could be impacted by loss of parkin. Immunoblotting analysis of mitochondrial fractions after 3 days of paraquat exposure did not result in dramatic differences in pS65-Ub levels either upon loss of parkin (*park*^−/−^) or transgenic overexpression (*daG4*>*UAS-park*) (Supplementary Figure 1C). We therefore devised a paraquat pulse-chase assay to probe the effect of loss of parkin on pS65-Ub dynamics. Flies were exposed to paraquat for one day and then kept under normal conditions (i.e., no paraquat) before being analysed for pS65-Ub levels by immunoblotting. At early time points, pS65-Ub levels were reduced in *park*^−/−^ mitochondria compared with mitochondria from wild-type animals (Figure 2C, D), consistent with a diminished feed-forward cycle of Pink1-parkin-dependent pS65-Ub production as previously described (Ordureau et al., 2014). In contrast, at later time points, pS65-Ub levels were elevated in *park*^−/−^ mitochondria compared with wild-type flies, which likely reflects a defect in turnover of damaged mitochondria (Figure 2C, D). It is therefore likely that parkin participates in the feed-forward cycle to promote further parkin recruitment to damaged mitochondria, but is not strictly required for the production of pS65-Ub on mitochondria in response to paraquat.

We next sought to interrogate the pattern of paraquat-stimulated mitochondrial ubiquitination in the presence and absence of Pink1 or parkin. Analysing the four Ub chain types (linked at K6, K11, K48 and K63) that are most abundant on depolarised mitochondria and produced by Parkin *in vitro* (Ordureau et al., 2014), the relative proportions of all four chain types were unchanged in mitochondria from *Pink1*^−^ and *park*^−/−^ flies compared to wild-type animals in basal conditions (Figure 2E-H). In response to paraquat, only K6 chains increased in abundance on wild-type mitochondria, while K11 chains remained unchanged. Surprisingly, K48 and K63 chains decreased as a proportion of the total mitochondrial Ub, presumably due to a more substantial increase in monoubiquitination as previously described (Swatek et al., 2019). The paraquat-induced increase in K6 chains appeared to depend on Pink1 and parkin, although we note a trend towards increasing K6 levels in these mutants (Figure 2E). Our results are therefore consistent with other reports that the molecular function of parkin, rather than to amplify pS65-Ub, may be to produce either K6 chains or another Ub signal on the OMM following recruitment to damaged mitochondria by binding to pS65-Ub (Ordureau et al., 2015; Gersch et al., 2017).

### *parkin*-null flies have elevated basal pS65-Ub

In our initial pipeline for detection of pS65-Ub, we enriched mitochondria by differential centrifugation. The pS65-Ub levels of untreated *park*^−/−^ flies were not substantially elevated in these fractions, as determined by mass spectrometry (Figure 2B) and immunoblotting (Supplementary Figure 1C). However, when we analysed whole cell lysates, untreated *park*^−/−^ animals displayed a striking abundance of pS65-Ub that was readily detectable by immunoblotting (Figure 3A). We confirmed that this signal represented pS65-Ub as it was sensitive to treatment with the deubiquitinase USP2 (Supplementary Figure 1D). The effect was also observed upon ubiquitous knockdown of *parkin* (*daG4>UAS-park RNAi*), confirming the specificity of the effect for loss of parkin (Figure 3B). The inducible RNAi line allowed us to assess the tissue distribution of the pS65-Ub in these flies using tissue-specific drivers. Interestingly, here we found that the majority of the pS65-Ub originated from the muscle rather than neurons (Figure 3B). This contrasts with the pS65-Ub produced upon response to paraquat in wild-type flies, where it was enriched in heads (Figure 1E). However, when we enriched for neural tissues by harvesting heads, some pS65-Ub was detectable in *park*^−/−^ flies (Figure 3C).

**Figure 3:**
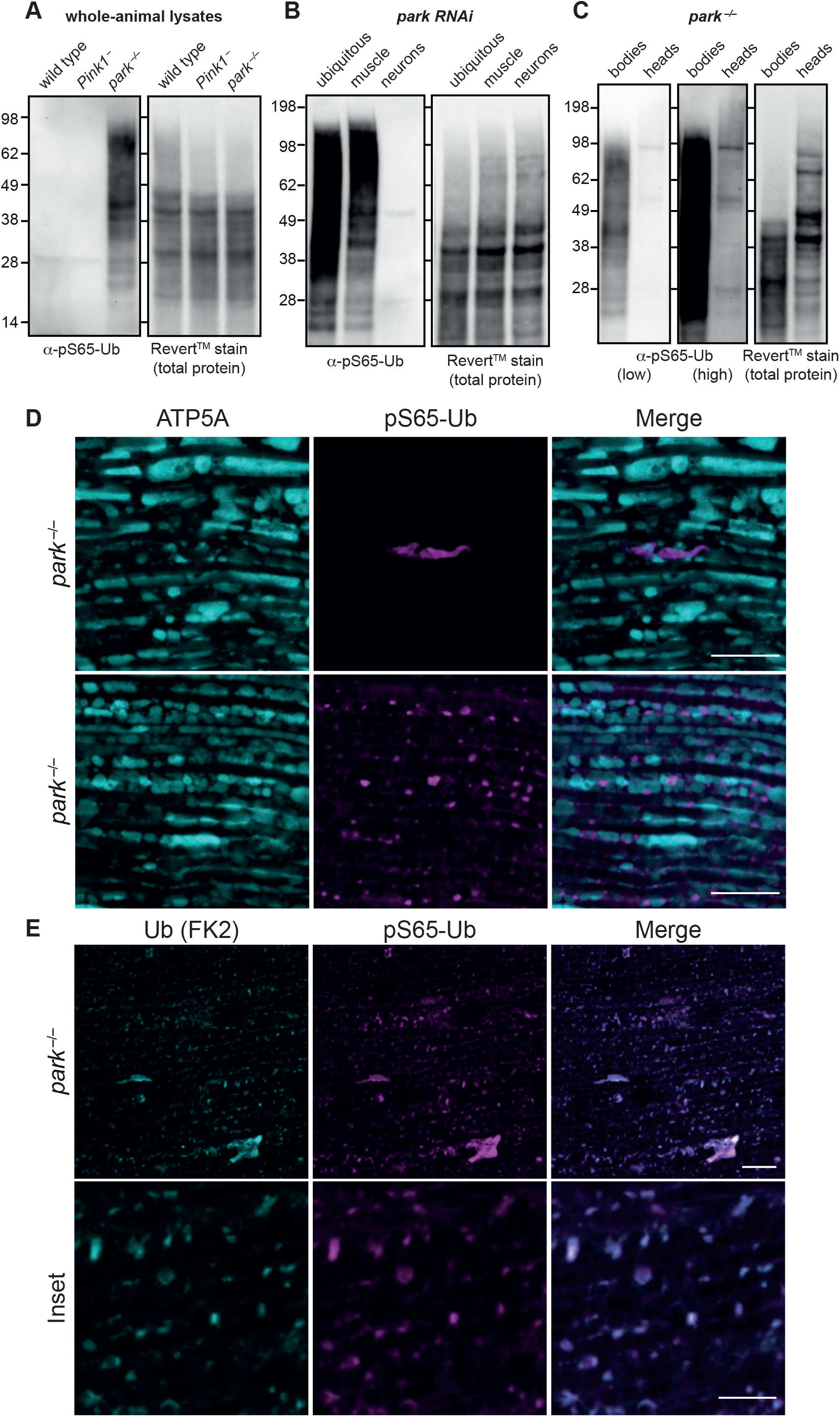
Loss of parkin affects both production and turnover of pS65-Ub. (A-B) pS65-Ub immunoblot of whole-animal lysates from untreated flies of the indicated genotypes. *park* RNAi was induced by *da-GAL4* (ubiquitous), *Mef2-GAL4* (muscle) or *nSyb-GAL4* (neurons). (C) pS65-Ub immunoblot of lysates from bodies or heads (as indicated) from *park*^−/−^ animals. (D) Flight muscles from approximately 3-day-old untreated *park*^−/−^ flies immunostained for ATP5A (IMM) and pS65-Ub. Scale bars = 10 μm. (E) Flight muscles from approximately 3-day-old untreated *park*^−/−^ flies immunostained for conjugated Ub (FK2) and pS65-Ub. Scale bars = 10 μm (top) and 5 μm (bottom).

To better understand the subcellular localisation of pS65-Ub in *park*^−/−^ flies we initially performed biochemical fractionation experiments. These results suggested that pS65-Ub localises to cellular membranes as opposed to the cytosol in *park*^−/−^ flies (Supplementary Figure 1D). We next employed an immunostaining approach, and noted the formation of heterogeneous pS65-Ub-positive structures in the flight muscles of *park*^−/−^ flies that were absent in wild-type and *Pink1*^−^ flight muscles (Figure 3D, Supplementary Figure 2A). These ranged from small punctate structures (<1 μm^3^) to very large objects that resembled hyperfused mitochondria but were mostly depleted for the mitochondrial marker ATP5A (Figure 3D). pS65-Ub staining appeared to show a greater degree of colocalization with an OMM-GFP marker than the inner mitochondrial membrane (IMM) protein ATP5A (Figure 3D, Supplementary Figure 2B). These structures clearly colocalised with a total Ub marker (Figure 3E) and were absent in *Pink1*^−^ flight muscles despite the presence of similar structures that stained for total Ub (Supplementary Figure 2C), confirming them as *bone fide* pS65-Ub. These results suggest that pS65-Ub accumulates on the OMM of dysfunctional mitochondria in the absence of parkin.

### Loss of core autophagy genes minimally affects pS65-Ub accumulation

The striking increase in pS65-Ub levels in *park*^−/−^ flies (Figure 3) indicated that turnover of pS65-Ub was disrupted, which presents a paradigm to investigate the turnover mechanisms downstream of Pink1 and parkin. Given the abundant evidence in cell culture models that PINK1-Parkin mediated turnover occurs via the canonical autophagy machinery (Lazarou et al., 2015; Nguyen et al., 2016), we analysed pS65-Ub levels in mutants of core autophagy genes, *Atg1* (homologue of ULK1), *Atg5* and *Atg8a* (homologue of LC3/GABARAP). We saw a modest age-related increase in pS65-Ub levels in *Atg5*^−^ (*Atg5*^5cc5^) flies compared with wild-type animals (Supplementary Figure 3A), but surprisingly, this was very low compared to the increase in pS65-Ub levels observed in *park*^−/−^ flies (Figure 4A). Consistent with this, neither loss of *Atg1* (*daG4>Atg1 RNAi*) nor *Atg8a*^−^ (*Atg8a*^KG07569^) led to the same dramatic increase in pS65-Ub levels as loss of *park* (Figure 4B, Supplementary Figure 3B).

**Figure 4:**
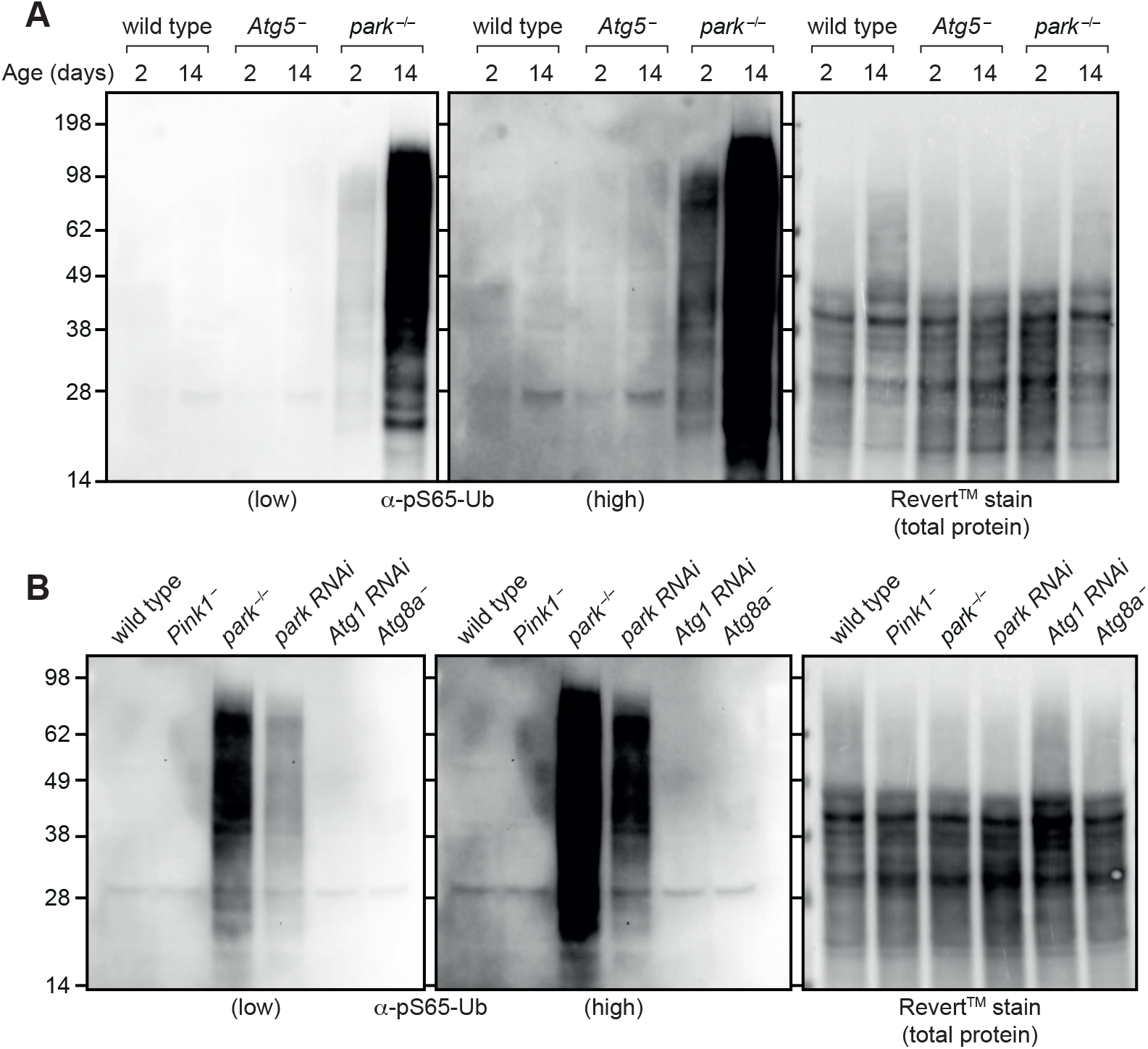
Loss of Atgs does not lead to the same degree of pS65-Ub accumulation as loss of parkin. (A) pS65-Ub immunoblot of whole-animal lysates from wild-type, *Atg5*^−^ and *park*^−/−^ animals harvested at the indicated ages. (B) pS65-Ub immunoblot of whole-animal lysates from young animals of the following genotypes: wild-type, *Pink1*^−^, *park*^−/−^, *park RNAi* (*daG4>UAS-park RNAi*), *Atg1 RNAi* (*daG4>UAS-Atg1 RNAi*), *Atg8a*^−^.

Loss of parkin led to pS65-Ub production that was readily detectable by immunoblotting as early as the larval stage of development (Supplementary Figure 3B). To further probe whether the canonical autophagy machinery affects pS65-Ub production, we quantified the size and number of pS65-Ub-positive puncta in larval muscle. Wild-type and *Pink1*^−^ larvae displayed no pS65-Ub puncta (Figure 5A, B, H), consistent with the absence of pS65-Ub observed by immunoblotting (Supplementary Figure 3B). In contrast, *park*^−/−^ tissues displayed abundant pS65-Ub puncta (Figure 5C, H). *Atg5*^−^ and *Atg8a*^−^ larvae also displayed pS65-Ub puncta, although they were markedly fewer and generally smaller than those present upon loss of parkin (Figure 5D, E, H, I), while *Atg5*^−^; *park*^−/−^ and *Atg8a*^−^; *park*^−/−^ double mutants displayed puncta similar in number and size to *park*^−/−^ alone (Figure 5F-I). These results suggest that canonical autophagy minimally contributes to the turnover of pS65-Ub-positive structures. Notably, while *park*^−/−^, *Atg5*^−^ and *Atg8a*^−^ animals are viable to adult stage, *Atg5*^−^; *park*^−/−^ and *Atg8a*^−^; *park*^−/−^ double mutants are generally non-viable past the pupal stage, with only a few rare escapers, indicating synthetic lethality from the combined effect of independent pathways.

**Figure 5:**
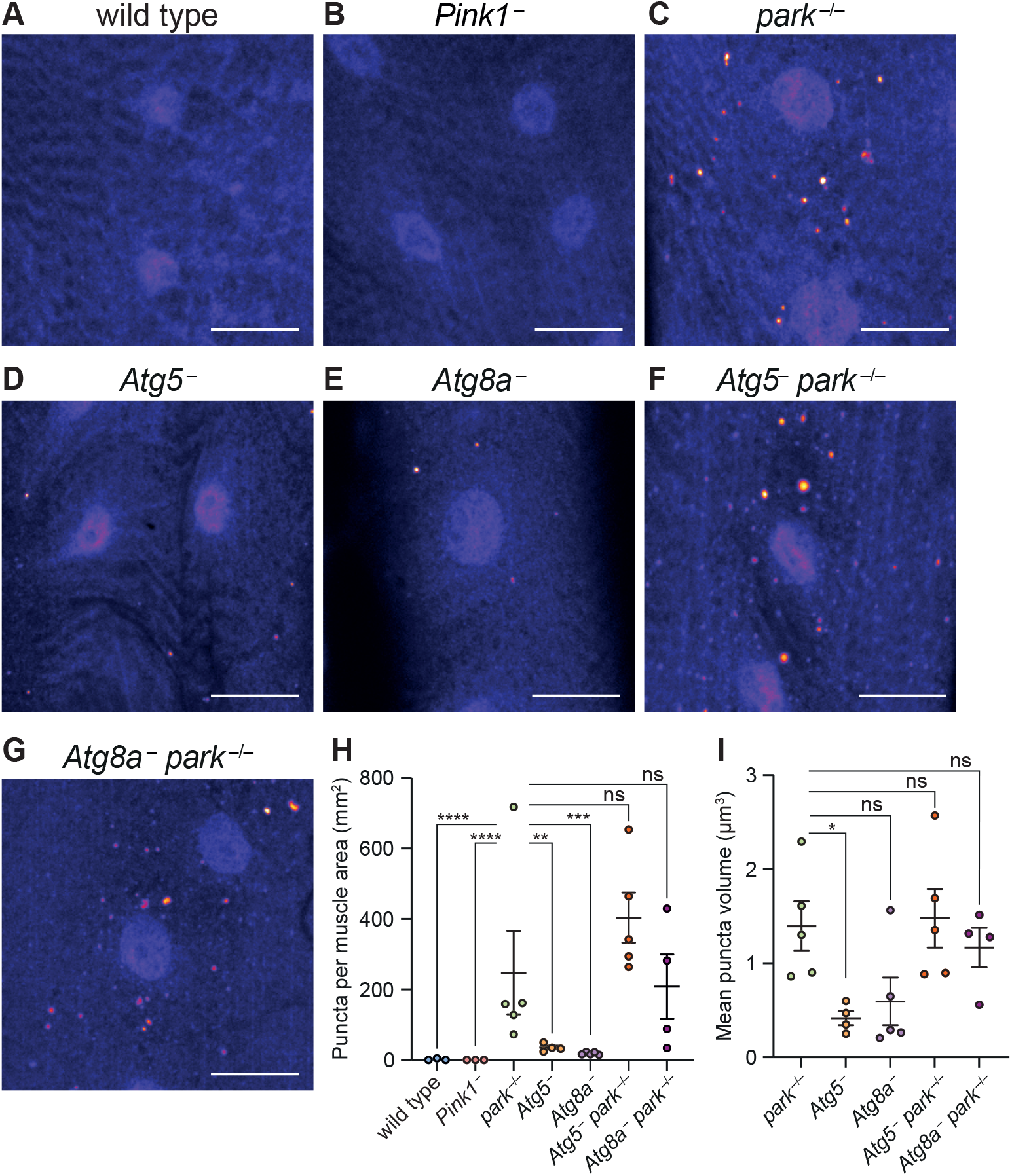
pS65-Ub immunostaining of larval muscle. (A-G) Representative images of pS65-Ub immunostaining false-coloured for intensity (Fire LUT) in muscle segments 6-7 from wandering L3 larvae of the following genotypes: (A) wild type, (B) *Pink1*^−^, (C) *park*^−/−^, (D) *Atg5*^−^, (E) *Atg8a*^−^, (F) *Atg5*^−^; *park*^−/−^, (G) *Atg8a*^−^; *park*^−/−^. Scale bar = 20 μm. (H) Quantification of number of pS65-Ub puncta from A-G displaying mean +/− SEM from the indicated number of animals (n = 3-5 as indicated). (I) Puncta volume from C-G expressed as mean +/− SEM (n = 4-5 animals). Statistical analysis used one-way ANOVA with Dunnett’s correction for multiple comparisons. * *P* < 0.05; ** *P* < 0.01; *** *P* < 0.001; **** *P* < 0.0001, ns = non-significant.

### parkin overexpression reduces pS65-Ub levels in the absence of Atg5

Although loss of the core autophagy components Atg1, Atg5 and Atg8a did not result in the same extent of pS65-Ub accumulation as loss of parkin, loss of Atg5 or Atg8a did lead to modestly increased pS65-Ub levels compared with wild-type animals (Supplementary Figure 3A, and Figure 5H). In addition, in the paraquat pulse-chase assay *Atg5*^−^ flies had elevated pS65-Ub at later time points relative to wild-type flies, suggestive of a block in turnover (Supplementary Figure 3C, D). These results are consistent with the autophagy machinery contributing to turnover of damaged mitochondria.

In order to further dissect whether the parkin-mediated pS65-Ub turnover is autophagy-dependent, we investigated the effect of parkin overexpression in an *Atg5*-null background (*Atg5*^5cc5^; *daG4>UAS-park*). We hypothesised that, if the Pink1-parkin pathway proceeds primarily via autophagy, then parkin overexpression should either not affect or perhaps even further increase pS65-Ub levels in an *Atg5*^−^ background. In contrast, if parkin drives autophagy-independent turnover, its overexpression should reduce pS65-Ub levels even in the absence of Atg5. We found in the paraquat pulse-chase assay that parkin overexpression substantially reduced pS65-Ub levels in an *Atg5*^−^ background relative to an *Atg5*^−^ mutant control (*Atg5*^5cc5^; *daG4>UAS-mito*-*HA*-*GFP*) (Figure 6A, B). We further confirmed by mass spectrometry that while mitochondria from *Atg5*^−^ flies displayed modestly elevated pS65-Ub levels, this could be reduced upon overexpression of parkin (Figure 6C). Taken together, these results indicate that parkin is able to drive pS65-Ub turnover independently of the canonical autophagy machinery in *Drosophila*.

**Figure 6:**
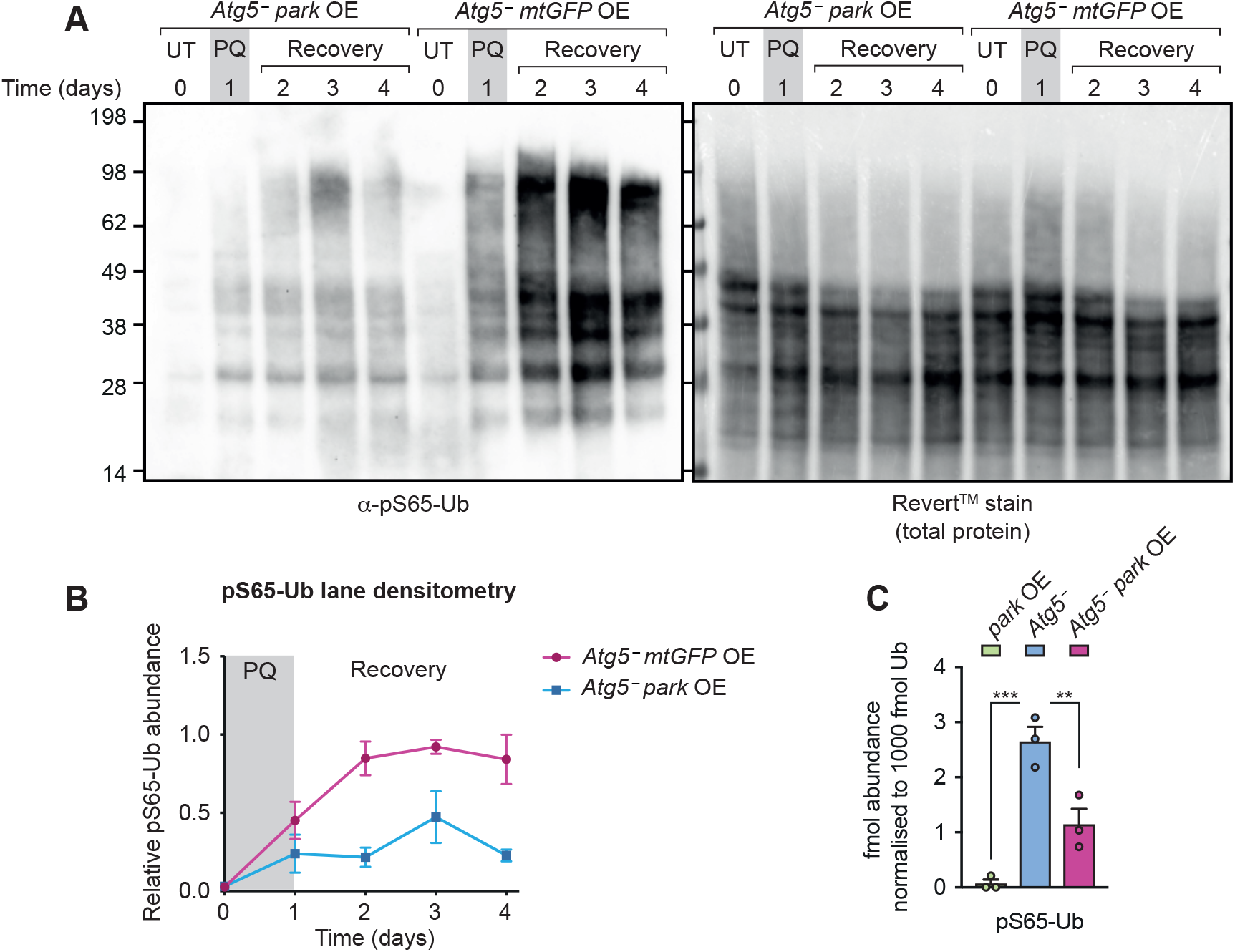
Parkin overexpression reduces pS65-Ub levels in an *Atg5*^−^ background. (A) pS65-Ub immunoblot of whole-animal lysates following paraquat pulse-chase assay of *Atg5*^−^ flies overexpressing parkin (*Atg5*^5cc5^ *daG4>UAS-park*) compared with *Atg5*^−^ flies overexpressing mitochondrially targeted GFP (*Atg5*^5cc5^ *daG4>UAS-mito*-*HA*-*GFP*). UT, untreated flies; PQ, 1-day paraquat treatment; Recovery, flies removed from paraquat and returned to normal food. (B) pS65-Ub lane densitometry, normalised to total protein (Revert^TM^ stain), of n = 3 independent replicates of (A), expressed as pS65-Ub intensity relative to the most intense band in each blot. Charts show mean +/− SEM. (C) Mass spectrometry analysis of normalised pS65-Ub levels in Ub-Clippase-treated, TUBE-enriched mitochondrial fractions from flies overexpressing parkin (*daG4>UAS-park*), *Atg5*^−^ flies and *Atg5*^−^ flies overexpressing parkin (*Atg5*^5cc5^ *daG4>UAS-park*). Charts show mean +/− SEM. Statistical analysis used One-way ANOVA with Dunnett’s correction for multiple comparisons. ** *P* < 0.01; *** *P* < 0.001.

## Discussion

We have optimised mass spectrometry and immunodetection methods to monitor physiological levels of pS65-Ub as a direct and specific readout of Pink1 activity in *Drosophila*, a preeminent model for dissecting the conserved functions of Pink1 and parkin. We have found that pS65-Ub is produced by Pink1 under basal conditions, albeit at very low levels. Our methods revealed that ~0.5% of total Ub on mitochondria from aged flies is Ser65-phosphorylated, and we further observed individual mitochondria that were enveloped in pS65-Ub. Although we could not reliably detect pS65-Ub in young flies without additional enrichment, we surmise that it is likely less than 0.1% of mitochondrial Ub. This suggests that under normal healthy conditions, in the absence of exogenous or accumulated endogenous stresses, Pink1 activation is either an extremely infrequent event or the pS65-Ub is very quickly degraded. This goes some way to explain why there was such negligible impact of loss of Pink1 or parkin on mitophagy reporters (Lee et al., 2018; McWilliams et al., 2018).

We also discovered that loss of parkin in *Drosophila* led to a striking increase in pS65-Ub levels in the absence of exogenous stimulation of the pathway. This surprising finding indicates that pS65-Ub alone is insufficient to elicit mitochondrial turnover, and therefore that parkin’s molecular function is not simply to amplify the pS65-Ub signal produced by Pink1. Several earlier studies also support this conclusion: increased stoichiometry of mitochondrial Ub phosphorylation was found to be inhibitory to mitophagy receptor recruitment in cell culture studies (Ordureau et al., 2018), and *in vitro* binding studies using Ser65-phosphorylated Ub chains have found that Ub phosphorylation does not promote autophagy receptor binding (Ordureau et al., 2015; Heo et al., 2015). Indeed, to our knowledge, Parkin itself is the only protein that has been shown to bind preferentially to pS65-Ub (Wauer et al., 2015). Our findings therefore suggest that the function of pS65-Ub is primarily to recruit parkin to damaged mitochondria, rather than to promote downstream organelle turnover.

What then is the molecular role of parkin? This study did not investigate the specific substrates that are ubiquitinated by parkin, as previous studies have found a breadth of OMM substrates ubiquitinated *in vivo* and in cell culture studies in a parkin-dependent manner (Martinez et al., 2017; Ordureau et al., 2018, 2020). We did, however, investigate the four main Ub chain types that are known to be produced by parkin – K6, K11, K48 and K63 (Ordureau et al., 2014) – and found that K6 chains, but not any other chain type, increased in relative abundance in mitochondrial fractions upon exposure to paraquat in a manner that was dependent on both Pink1 and parkin. This result was particularly interesting given that the only mitochondria-resident deubiquitinase, USP30, preferentially binds K6 chains (Cunningham et al., 2015; Wauer et al., 2015; Gersch et al., 2017; Sato et al., 2017). It is therefore possible that the primary function of parkin on mitochondria is to produce K6 chains. However, the functions of K6 chains are not fully understood (Swatek and Komander, 2016); further work is required to elucidate the precise contribution of this atypical Ub chain type to mitochondrial quality control.

The dramatic increase in pS65-Ub levels upon loss of parkin allowed us to assess the machinery responsible for downstream turnover of pS65-ubiquitinated mitochondria. Analysing the autophagy machinery, we observed that upon loss of the core autophagy components Atg1, Atg5 and Atg8a, pS65-Ub levels were not affected nearly to the extent observed in *park*^−/−^ flies, which suggested that pS65-Ub is not primarily turned over via canonical autophagy. Moreover, parkin overexpression was able to reduce both basal and paraquat-induced pS65-Ub levels in an *Atg5*^−^ background. These results add to the growing evidence that the Pink1-parkin pathway may, under more physiological conditions, promote turnover of damaged mitochondrial components in an autophagy-independent manner. For instance, Vincow et al. analysed turnover rates of mitochondrial proteins in *Drosophila* and found that Pink1 and parkin were required for the turnover of a subset of IMM proteins that was distinct from those turned over by the core autophagy protein Atg7 (Vincow et al., 2013). The authors also found some overlap between the proteins turned over by parkin and Atg7, and we observed a slight accumulation of pS65-Ub in autophagy-deficient flies, consistent with Pink1-parkin mediated degradation occurring partially via a classic autophagy route. However, the Pink1-parkin pathway clearly has roles that are divergent from canonical autophagy, as *Atg5*^−^; *park*^−/−^ and *Atg8a*^−^; *park*^−/−^ double mutants showed synthetic lethality, and parkin overexpression was able to reduce both basal and paraquat-induced pS65-Ub levels in an *Atg5*^−^ background. Consistent with this, the ability of parkin overexpression to rescue *Pink1* phenotypes has been shown to be unaffected by loss of Atg1 or Atg18 (Liu and Lu, 2010).

One alternative mechanism of pS65-Ub turnover that is promoted by parkin could be direct proteasomal degradation of pS65-ubiquitinated OMM proteins (Tanaka et al., 2010; McLelland et al., 2018). Alternatively, growing evidence points to the existence of MDVs, defined as small (~100 nm) cargo-selective vesicles that form independently of the autophagy machinery (Sugiura et al., 2014). Multiple studies have described a role for Pink1 and parkin in the formation of a subset of these vesicles that are delivered to lysosomes (McLelland et al., 2014; Ryan et al., 2020). However, *in vivo* evidence for the existence of MDVs is limited, due in part to technical constraints in observing such small structures in complex tissues. We anticipate that the methods described herein could aid in validating the presence of MDVs and their dependence on Pink1 and parkin *in vivo*, or conversely whether parkin primarily drives turnover of pS65-ubiquitinated proteins in a proteasome-dependent manner.

### pS65-Ub as a biomarker for neurodegeneration

Exposure to sub-lethal doses of paraquat led to a strong induction of pS65-Ub production in *Drosophila*. This result supports a link between PD caused by environmental exposure to mitochondrial toxins (Tanner et al., 2011) and genetic parkinsonism caused by loss of *PINK1* and *PRKN* gene function (Valente et al., 2004; Kitada et al., 1998). pS65-Ub has been proposed as a potential biomarker for neurodegenerative disease, with a recent study finding elevated pS65-Ub levels in blood samples from a cohort of Alzheimer’s Disease patients compared with age-matched controls (Watzlawik et al., 2020). The cellular pathology underlying PD precedes classical symptom onset and therefore clinical diagnosis by many years, which is likely to hamper the success of clinical trials of potentially disease-modifying drugs as they may be given too late to halt disease progression (Stern et al., 2012). It is therefore of vital importance to identify patients as early in their disease progression as possible. We found that pS65-Ub was readily detectable in *park*^−/−^ animals in developmental stages prior to overt neurodegeneration, thereby suggesting that pS65-Ub accumulation is an early event that, if replicated in mammals, could have potential as an early-stage diagnostic biomarker. However, pS65-Ub levels increased with healthy ageing and was absent upon loss of Pink1, so we posit that a healthy range of pS65-Ub abundance would need to be established in order for pS65-Ub levels to be useful as a clinical PD diagnostic tool.

## Conclusions

We have developed methods to detect pS65-Ub at physiological levels in *Drosophila*, and delivered the unanticipated finding that loss of parkin results in a striking elevation in pS65-Ub levels that is not recapitulated upon loss of the canonical autophagy genes *Atg5*, *Atg8a* or *Atg1*. We expect that the tools described herein will greatly aid in future studies dissecting the downstream mechanisms of mitochondrial turnover that are promoted by Pink1 and parkin *in vivo*.

## Materials and Methods

### *Drosophila* stocks and husbandry

Flies were raised under standard conditions in a temperature-controlled incubator with a 12h:12h light:dark cycle at 25 °C and 65 % relative humidity, on food consisting of agar, cornmeal, molasses, propionic acid and yeast. The following strains were obtained from the Bloomington *Drosophila* Stock Centre (RRID:SCR_006457): *w*^1118^ (RRID: BDSC_6326), *da*-*GAL4* (RRID: BDSC_55850), *Mef2*-*GAL4* (RRID: BDSC_27390), *nSyb*-*GAL4* (RRID: BDSC_51941), *Atg8a*^KG07569^ (RRID: BDSC_14639), *UAS-mito*-*HA*-*GFP* (RRID: BDSC_8443) and *UAS-park RNAi* (RRID: BDSC_38333). The *UAS-Atg1 RNAi* (VDRC_16133) line was obtained from the Vienna *Drosophila* Resource Centre. *Pink1*^B9^ flies were a kind gift of J. Chung (Park et al., 2006), and the *Atg5*^5cc5^ stock was a kind gift of G. Juhasz (Kim et al., 2016). *UAS-park*C2, *park*^25^, *UAS-mito*-*APOBEC1* and *UAS-mitoQC* lines have been described previously (Greene et al., 2003; Andreazza et al., 2019; Lee et al., 2018). Male flies only were used for experiments in adults, while experiments in larvae used both males and females except animals with X chromosome balancers (*Pink1*^B9^, *Atg5*^5cc5^), for which only male animals were used. For ageing experiments, flies were maintained in bottles (MS experiments) or tubes (immunostaining experiments), transferred to fresh food thrice weekly, and harvested after 50 to 60 days.

### Paraquat exposure assays

For MS experiments, flies were maintained in bottles (100 to 200 flies per replicate) containing 9 semi-circular pieces of filter paper [90 mm diameter, Cat ID. 1001-090] saturated with 5 % (w/v) sucrose solution containing 5 mM paraquat. Sucrose-only starvation experiments were performed as above, with the omission of paraquat. After 3 days, the flies were anaesthetised with mild CO_2_ and live flies only were harvested. For pulse-chase experiments, 5-25 flies were harvested per replicate. The day 0 control was taken prior to paraquat treatment, and the remaining flies were incubated overnight in bottles containing paraquat as above. The next day, flies were anaesthetised with CO_2_, dead flies were removed, a day 1 timepoint was taken, and the remaining flies were divided among tubes of food with no more than 20 flies per tube. At day 2, all flies were flipped onto fresh food, and then flipped every 2-3 days thereafter until harvest.

### Mitochondrial enrichment by differential centrifugation

All steps were performed on ice or at 4 °C. For mass spectrometry analysis, mitochondria were harvested from fresh (not frozen) flies according to (Lazarou et al., 2007), with modifications. Whole flies (60-200 per replicate) were placed in a dounce homogeniser, Solution A (70 mM sucrose, 20 mM HEPES pH 7.6, 220 mM mannitol, 1 mM EDTA) containing cOmplete protease inhibitors (Roche) and PhosSTOP phosphatase inhibitors (Roche) was added (approximately 10 μL per fly), and the flies were homogenised with 35 strokes of a drill-fitted pestle. The homogenate was transferred to a 50 mL tube and incubated 30 minutes, then centrifuged for 5 minutes at 1,000 x g. The supernatant (containing mitochondria) was transferred to microcentrifuge tubes and centrifuged 15 minutes at 10,000 x g. The post-mitochondrial supernatant was removed and the pellet (containing mitochondria) was resuspended in Solution A. The homogenate was then clarified by centrifugation for 5 minutes at 800 x g, and the supernatant transferred to a fresh tube. This clarification step was repeated once more to ensure all cuticle was removed from the sample. The supernatant was then centrifuged 10 minutes at 10,000 x g, and the post-mitochondrial supernatant was discarded. The pellet was resuspended in Solution A and centrifuged 10 minutes at 10,000 x g, and this wash step was repeated for a total of three times. The washed pellet was resuspended in Sucrose Storage Buffer (500 mM sucrose, 10 mM HEPES pH 7.6) and stored at −80 °C until needed.

For immunoblotting analysis and biochemical fractionation from small numbers of flies (10-30), a modified mitochondrial enrichment procedure was performed. Flies were prepared either fresh or after flash-freezing in liquid nitrogen, with all direct comparisons performed with flies that were prepared in the same manner. Flies were transferred into a Dounce homogeniser containing 700 μL Solution A containing protease and phosphatase inhibitors as above, and manually homogenised with 50 strokes of a pestle. The homogenate was transferred to an Eppendorf tube, a further 500 μL of Solution A was added to the homogeniser and the sample was homogenised with a further 10 strokes. The homogenates were pooled and incubated for 30 minutes, then centrifuged for 5 minutes at 800 x g. The supernatant (containing mitochondria) was transferred to a new tube and clarified twice by centrifugation for 5 minutes at 1,000 x g. The clarified supernatant was then centrifuged for 10 minutes at 10,000 x g and the post-mitochondrial supernatant was discarded or, in the case of biochemical fractionation experiments, further centrifuged for 30 minutes at 21,000 x g, and the pellet and supernatant retained for analysis. The mitochondrial pellet was washed once in Solution A containing only protease inhibitors, and then once in Solution A without inhibitors. The washed mitochondrial pellet was resuspended in 50 to 200 μL Sucrose Storage Buffer, the protein content determined by BCA assay (Thermo Pierce), and stored at −80 °C until needed.

### USP2 treatment

For the validation of pS65-Ub signal in *park*^−/−^ samples, 30 μg protein per subcellular fraction was treated with the pan-specific deubiquitinase USP2 (BostonBiochem, E-506). The USP2 enzyme was diluted in buffer (50 mM Tris pH 7.5, 50 mM NaCl, 10 mM DTT) (Hospenthal et al., 2015) and then added to the subcellular fractions to a final USP2 concentration of 1 μM. The mixture was incubated for 45 minutes at 37 °C prior to analysis by immunoblotting.

### Mass spectrometry sample preparation and analysis

Absolute quantification (AQUA) analysis of Ub modifications was performed using Ub-Clipping (Swatek et al., 2019), with modifications. 500 μg mitochondria (from approximately 100 flies) prepared as above were resuspended in 250 μL TUBE lysis buffer (PBS containing 1% (v/v) NP-40, 2 mM EDTA, 10 mM chloroacetamide, cOmplete EDTA-free protease inhibitor cocktail (Roche)) supplemented with 8 μg/ mL GST-Ubiquilin-UBA (Fiil et al., 2013; Hrdinka et al., 2016; Hjerpe et al., 2009). The lysate was incubated on ice for 20 minutes then centrifuged 15 minutes at 21,000 x g, 4 °C. The clarified lysate was added to 20 μL Glutathione Sepharose 4B resin (GE Healthcare) that had been washed three times in TUBE lysis buffer, and was incubated 2 h at 4 °C with gentle rotation. The lysate was removed and the beads were washed twice with PBS containing 0.1% (v/v) Tween 20, then twice with PBS. 80 μL Lb^pro^ reaction buffer (50 mM NaCl, 50 mM Tris pH 7.4, 10 mM DTT) containing 20 μM Ub-clippase (Lb^pro^ construct containing residues 29-195 with L102W mutation, purified as described previously (Swatek et al., 2019; Guarné et al., 2000)) was added and Ub was cleaved from the beads for 16 h at 37 °C. The supernatant was removed, the beads were washed with Lb^pro^ reaction buffer, and the supernatants pooled and acidified to pH <4 using formic acid (FA), prior to fractionation using StageTips (Rappsilber et al., 2007). StageTips were assembled using 4 plugs that were cut using a gauge 16 needle (Hamilton) from C_4_ substrate (SPE-Disks-Bio-C4-300.47.20, AffiniSEP) and assembled into a P200 pipette tip using a plunger (Hamilton). The matrix was activated by the addition of 30 μL methanol and the tip was centrifuged inside a 2 mL Eppendorf tube at 800 x g for 30 seconds at room temperature to allow the liquid to pass through. The tip was then equilibrated by passing through 30 μL 80% (v/v) acetonitrile (ACN), 0.1% (v/v) FA, followed by 30 μL 0.1% (v/v) FA. The acidified sample was loaded and centrifuged as above until almost all the liquid had passed through. The tip was then desalted by passing through 40 μL 0.1% (v/v) FA, twice. The StageTip was then washed twice with 30 μL 20% (v/v) ACN, 0.1% (v/v) FA, then the ubiquitin was eluted into a clean tube with two elutions of 30 μL 45% (v/v) ACN, 0.1% (v/v) FA. The eluate was lyophilised and resuspended in Trypsin Resuspension Buffer (Promega) supplemented with Tris pH 8.0 to ensure a final pH above 6. Sequencing grade modified Trypsin (Promega) was added at a concentration of 1 μg per 250 μg initial mitochondrial protein, and the samples were incubated 16 h at 37 °C. For StageTip purification after trypsin treatment, the sample was acidified to pH <4 using FA. AQUA peptides, supplied by Cambridge Research Biochemicals (pS65-Ub peptide) and Cell Signalling Technologies (all other peptides), were spiked in at concentrations as indicated in the Supplementary Table 1 and the sample was loaded into a StageTip containing 4 plugs of C_18_ substrate (SPE-Disks-Bio-C18-100.47.20, AffiniSEP) that had been assembled, activated and pre-equilibrated as above. The tip was washed 3 times in 0.1% (v/v) FA, then elution was performed twice with 30 μL 80% (v/v) ACN, 0.1% (v/v) FA. Samples were lyophilised and resuspended in 5% (v/v) ACN, 0.1% (v/v) FA, and 10μL was injected onto a Dionex Ultimate 3000 HPLC system (Thermo Fisher Scientific), and trapped on a C18Acclaim PepMap100 (5μm, 100μm x 20 mm nanoViper; Thermo Scientific). Peptides were eluted with a 60-minute acetonitrile gradient (2-40%) at a flow rate of 0.3 μL min^−1^. The analytical column outlet was directly interfaced via an EASY-Spray electrospray ionisation source to a Q Exactive mass spectrometer (Thermo Fisher Scientific). The following settings were used: resolution, 140,000; AGC target, 3E6; maximum injection time, 200 ms; scan range, 150-2,000 m/z. Absolute abundances of Ub peptides were calculated by peak integration using Xcalibur Qual Browser (Version 2.2, Thermo Fisher Scientific). Layouts were applied according to the Supplementary Table 1, and abundances were calculated relative to the known amount of added AQUA reference peptide using Microsoft Excel.

For the detection of pS65-Ub in young flies (Figure 1B), the following modifications to the method were performed. Instead of TUBE-mediated Ub pulldown, mitochondrial fractions were sodium carbonate-extracted to enrich ubiquitinated integral membrane proteins as previously described (Swatek et al., 2019). In brief, 4 mg mitochondria were resuspended in 4 mL 100 mM Na_2_CO_3_. The mixture was incubated 30 minutes on ice with occasional vortexing, then centrifuged 30 minutes, 21,000 x g, 4 °C. The supernatant, containing soluble and peripheral membrane proteins, was discarded and the pellet, containing integral membrane proteins, was then resuspended in Lb^pro^ reaction buffer (1 μL per 10 μg mitochondria). An equal volume of 20 μM Lb^pro^ was added (10 μM final concentration) and the mixture was incubated overnight at 37 °C. The samples were centrifuged 30 minutes, 21,000 x g, 4 °C, and the supernatant was acidified and purified using StageTips as above (1 StageTip per 1 mg starting material). Trypsin treatment was performed as above and AQUA peptides were spiked in according to the Supplementary Table 1. Phospho-peptide enrichment was then performed using the High-Select^TM^ TiO_2_ Phospho-peptide Enrichment kit (Thermo Fisher Scientific). Each replicate was divided between two TiO_2_ columns and prepared according to the manufacturer’s instructions. The eluates were pooled, lyophilised and analysed by LC-MS as above.

### Antibodies and dyes

The following mouse antibodies were used for immunoblotting (WB) and/or immunofluorescence (IF) in this study: ATP5A (ab14748, 1:10000 (WB), 1:300 (IF, adult muscle)), Ubiquitin (clone FK2, D058-3, 1:2000 (WB), 1:250 (IF, adult muscle), Actin (MAB1501, 1:1000 (WB)), GAPDH (GTX627408, 1:1000 (WB)). The following rabbit antibodies were used in this study: pS65-Ub (62802S, 1:750 (WB), 1:200 (IF, larval muscle), 1:120 (IF, adult muscle)), COXIV and SDHA (both kind gifts from Edward Owusu-Ansah (Murari et al., 2020), 1:2000 (WB)), Porin (PC548, 1:5000 (WB)). The following secondary antibodies were used: sheep anti-mouse (HRP-conjugated, NXA931V, 1:10000 (WB)), donkey anti-rabbit (HRP-conjugated, NA934V, 1:10000 (WB)), goat anti-mouse (AlexaFluor 488, A11001, 1:200 (IF)), goat anti-rabbit (AlexaFluor 594, A11012, 1:200 (IF)), goat anti-rabbit (AlexaFluor 647, A21244, 1:200 (IF)).

### Whole-animal lysis and immunoblotting

For the analysis of pS65-Ub levels in whole cell lysates by immunoblot, 180 μL cold RIPA buffer (150 mM NaCl, 1% (v/v) NP-40, 0.5 % (w/v) sodium deoxycholate, 0.1% (w/v) SDS, 50 mM Tris pH 7.4), supplemented with cOmplete and PhosSTOP inhibitors, was added to 2 mL tubes containing 1.4 mm ceramic beads (Fisherbrand 15555799). Animals (5 to 20 per replicate) were harvested and stored on ice or flash-frozen in liquid N_2_, with all direct comparisons performed with flies that were harvested in the same manner. The flies were added to the tubes containing RIPA buffer and lysed using a Minilys homogeniser (Bertin Instruments) with the following settings: maximum speed, 10 seconds on, 10 seconds on ice, for a total of three cycles. After lysis, samples were returned to ice for 10 minutes then centrifuged 5 minutes at 21,000 x g, 4 °C. 90 μL supernatant was transferred to a fresh Eppendorf tube and centrifuged a further 10 minutes at 21,000 x g. 50 μL supernatant was then transferred to a fresh Eppendorf tube and the protein content determined by BCA assay as above. 30 μg total protein was then diluted in NuPAGE LDS loading dye (Invitrogen) and analysed by SDS-PAGE using Novex 4-12% Bis-Tris NuPAGE gels (Invitrogen). For the analysis of mitochondria-enriched fractions, 30 to 50 μg mitochondrial protein was aliquoted into a tube, centrifuged 10 minutes at 16,000 x g, the supernatant removed and the pellet resuspended in LDS loading dye prior to SDS-PAGE analysis as above. Gels were transferred onto pre-cut and -soaked PVDF membranes (1704157, BioRad) using the BioRad Transblot Turbo transfer system, and blots were immediately stained with Revert total protein stain (LiCOR) where indicated, according to the manufacturer’s instructions. Fluorescence intensity was measured using a BioRad Chemidoc MP using the IR680 setting. Blots were then washed by gentle shaking 3 times for 5 minutes in PBS containing 0.1% (v/v) Tween-20 (PBST), and blocked by incubation with PBST containing 5% (w/v) skim milk for 30 minutes. Blots were washed a further 3 times as above then incubated at 4 °C overnight with primary antibodies in PBST containing 3 % (w/v) BSA. A further 3 washes were performed then the blots were incubated for one hour in secondary antibodies made up in PBST containing 5% (w/v) skim milk. Blots were then washed twice in PBST and once in PBS (twice in the case of pS65-Ub blots) prior to incubation with ECL reagent. For pS65-Ub blots, SuperSignal Femto reagent (Thermo Scientific) was used, while other blots used Clarity ECL reagent (BioRad). Blots were imaged using the BioRad Chemidoc MP using exposure settings to minimise overexposure, except where high exposure is indicated. Image analysis was performed using Image Lab (Version 5.2.1 build 11, BioRad) and images were exported as TIFF files for presentation.

### Immunostaining of *Drosophila* tissues

Larval filet and adult thoraces dissections were performed in PBS and fixed in 4% formaldehyde, pH 7.0, for 20 (larval filet) or 30 (adult flight muscles) minutes respectively. Permeabilisation was performed for 30 minutes in PBS containing 0.3% (v/v) Triton X-100 (PBS-TX), then tissues were blocked for 1 h in PBS-TX containing 1% (w/v) BSA. Primary antibody incubation was performed overnight at 4 °C in PBS-TX containing 1% (w/v) BSA. The tissues were washed 3 times for 10 minutes each in PBS-TX prior to incubation with secondary antibodies in PBS-TX containing 1% (w/v) BSA for 2 h at room temperature (larval filet) or overnight at 4 °C (adult thoraces). The tissues were washed three times for 10 minutes in PBS-TX, then once for 10 minutes in PBS, and rinsed once in water prior to mounting in Prolong Diamond Antifade mounting media with DAPI (Thermo Fisher Scientific).

### Microscopy and image analysis

Fluorescence microscopy imaging was performed using a Zeiss LSM 880 confocal microscope equipped with a 20x Plan Apochromat (air, NA = 0.8) and 63x Plan Apochromat (oil immersion, NA = 1.4) objective lenses. Laser power and gain settings were adjusted depending on the fluorophore, but were maintained across samples for the purpose of comparing pS65-Ub levels among genotypes. For imaging OMM-GFP, the GFP component of mito-QC constructs (UAS-mCherry-GFP-Fis1101-152) was imaged in conjunction with AlexaFluor 647 for pS65-Ub.

Images were processed using FIJI (ImageJ, Version 2.1.0/1.53c) for figure presentation. For the quantification of pS65-Ub puncta in larval muscle 6-7 (Figure 5H), z-stacks (5 per image, 0.28 μm step size) were cropped to remove extraneous tissue, axons, and neuromuscular junctions, which we observed contained high background signal with the anti-pS65-Ub antibody. The retained area was measured, local background subtraction was performed, and the number of puncta quantified using the 3D Object Counter v2.0 program on FIJI using the same threshold value for all samples. Puncta with a size smaller than 0.1 μm^3^ were excluded, and the remaining puncta considered to be true pS65-Ub puncta. The data were imported into Prism and a frequency distribution analysis performed to obtain number and mean volume of puncta.

### Statistical Analysis

Statistical analyses were performed using Prism (Version 9.1.0 (216)). For the analysis of mass spectrometry data presented in Figure 2, each Ub modification was analysed by Ordinary one-way ANOVA with Šidák’s correction for multiple comparisons (wild-type untreated compared with *Pink1*^−^ and *park*^−/−^ untreated, and +/− paraquat comparison within each genotype, five comparisons total). The pS65-Ub abundance presented in Figure 6C was analysed by Ordinary one-way ANOVA with Dunnett’s correction for multiple comparisons (each genotype compared with *Atg5*^−^, two comparisons total). For the analysis of number of pS65-Ub puncta in Figure 5H, data were first log-transformed (Y = Ln(y+1)) to account for heteroscedasticity in the raw data. The transformed data, as well as the raw data in Figure 5I, were analysed by One-way ANOVA with Dunnett’s multiple comparisons (each genotype compared with *park*^−/−^).

## Data Availability

All data generated or analysed during this study are included in the manuscript and supporting files.

## Author Contributions

JLU Conceptualization, Data curation, Formal analysis, Validation, Methodology, Writing - original draft.

JLL Formal analysis, Methodology, Writing - review and editing.

ASM Formal analysis, Methodology, Writing - review and editing.

AJW Conceptualization, Formal analysis, Supervision, Funding acquisition, Investigation, Project administration, Writing – original draft, review and editing.

## Acknowledgements

This work is supported by Medical Research Council core funding (MC_UU_00015/6). JLU was supported by a Gates Cambridge Scholarship. We thank Prof. D. Komander for reagents and for support in the early phase of this project. The funders had no role in study design, data collection and analysis, decision to publish, or preparation of the manuscript. Stocks were obtained from the Vienna *Drosophila* Stock Centre and the Bloomington *Drosophila* Stock Center which is supported by grant NIH P40OD018537. We thank Prof. G Juhasz for generously sharing fly stocks, and we thank Whitworth lab members for discussions and critical reading of the manuscript.

---

**Supplementary Figure 1:**
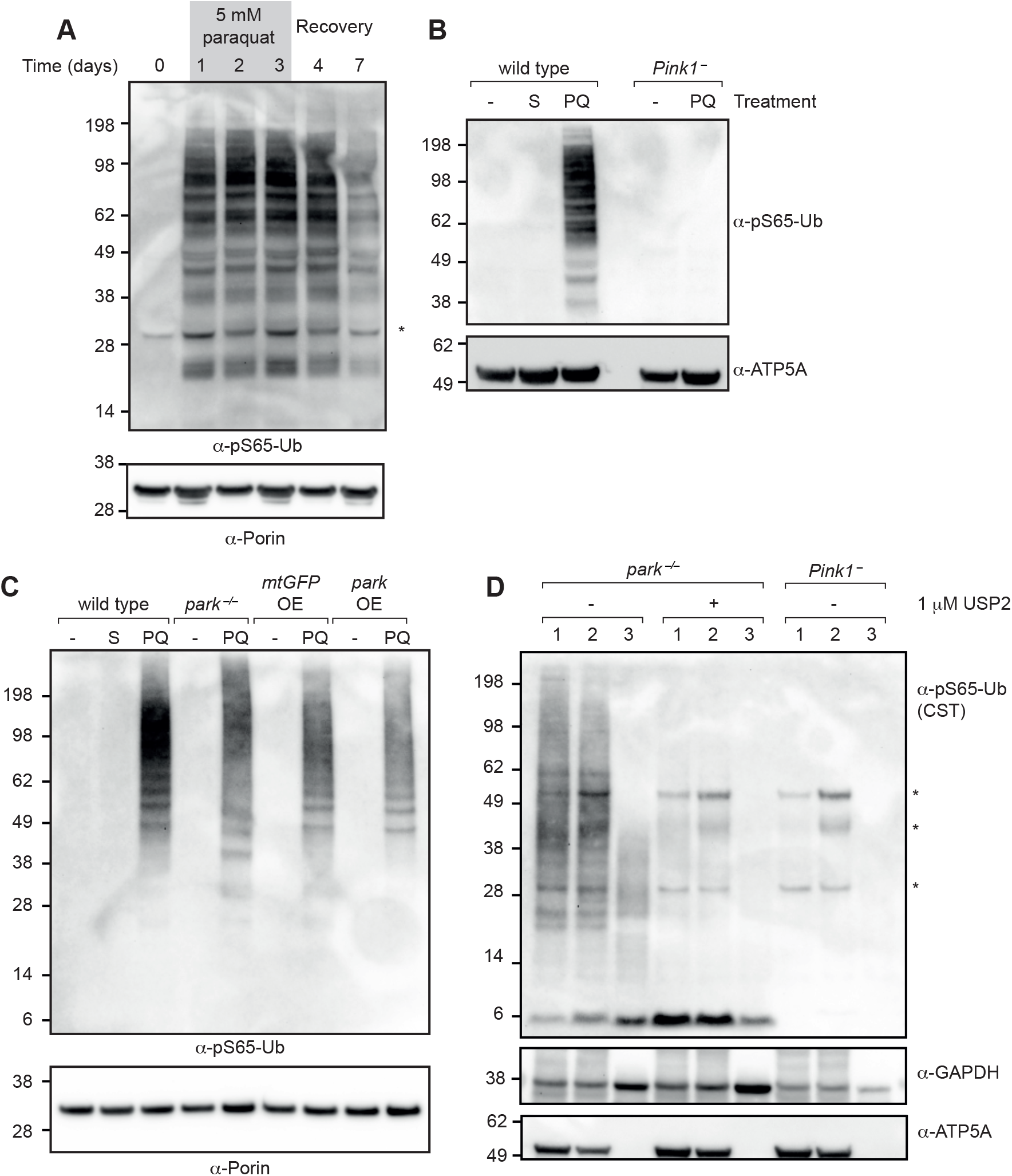
(A) pS65-Ub immunoblot of mitochondria-enriched fractions from wild-type flies treated for the indicated number of days with paraquat. Recovery, return to normal food. (B) pS65-Ub immunoblot of mitochondrial fractions from wild-type and *Pink1*^−^ flies following either no treatment (−) or after treatment for 3 days with sucrose (S) or paraquat (PQ). (C) pS65-Ub immunoblot of mitochondria-enriched fractions of the following genotypes: wild type, *park*^−/−^, mtGFP overexpression (*daG4>UAS-mito*-*HA*-*GFP*), parkin overexpression (*daG4>UAS-park*). Note that lanes 2 and 3 are shown in Figure 1C. (D) pS65-Ub immunoblot following subcellular fractionation and USP2 treatment as indicated. 1, 10,000 x g pellet; 2, 21,000 x g pellet; 3, 21,000 x g supernatant. * = non-specific bands.

**Supplementary Figure 2:**
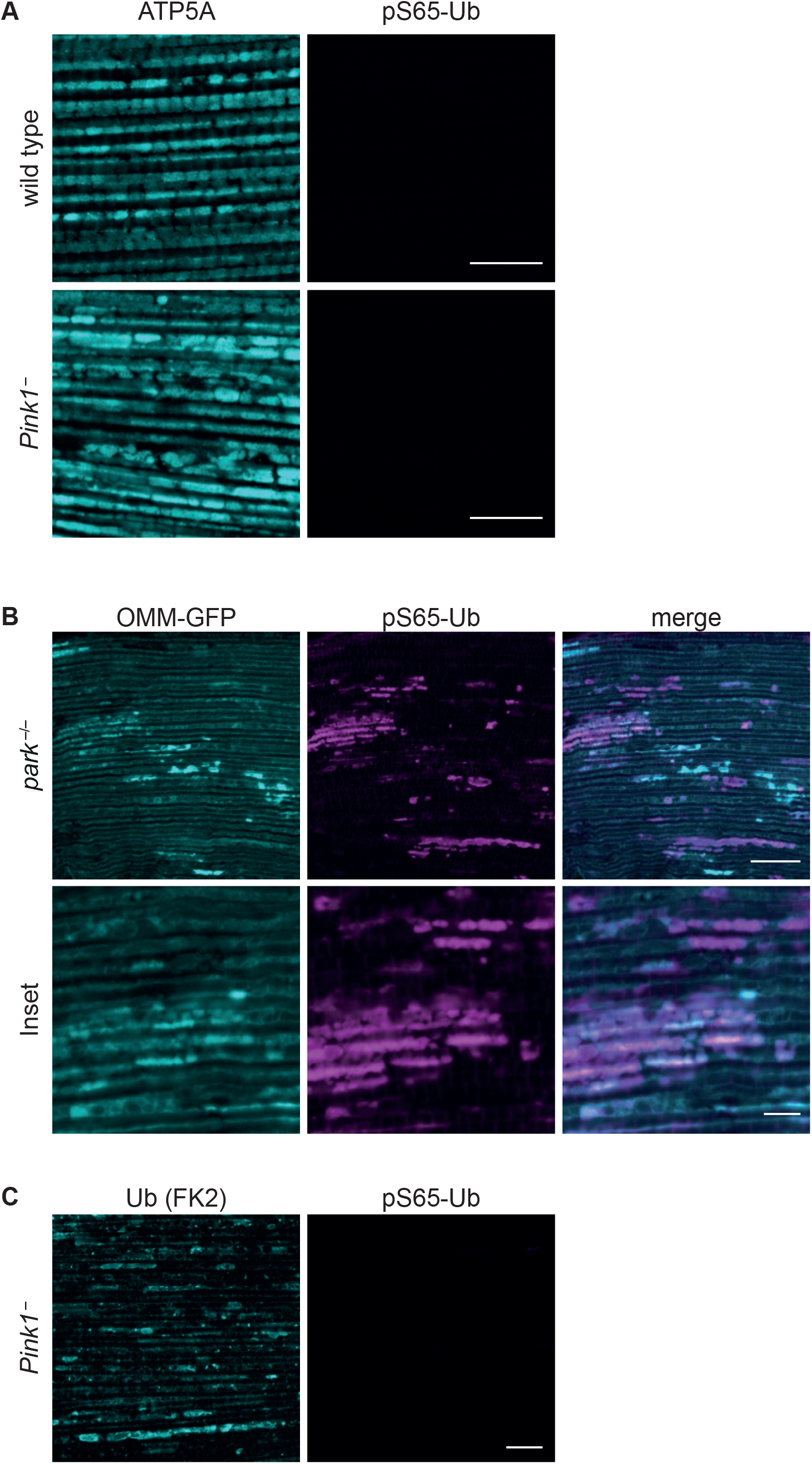
(A) Immunostaining of flight muscles of *Pink1*^−^ and wild-type flies. Note that signal acquisition, brightness and contrast settings for pS65-Ub are identical to those presented in Figure 3D. Scale bars = 10 μm. (B) pS65-Ub immunostaining (AlexaFluor 647) of flight muscles from *park*^−/−^ flies expressing the OMM-GFP marker. Scale bars = 20 μm (inset 5 μm). (C) Flight muscles from young *Pink1*^−^ flies, co-stained with Ub (FK2) and pS65-Ub. Scale bar = 10 μm.

**Supplementary Figure 3:**
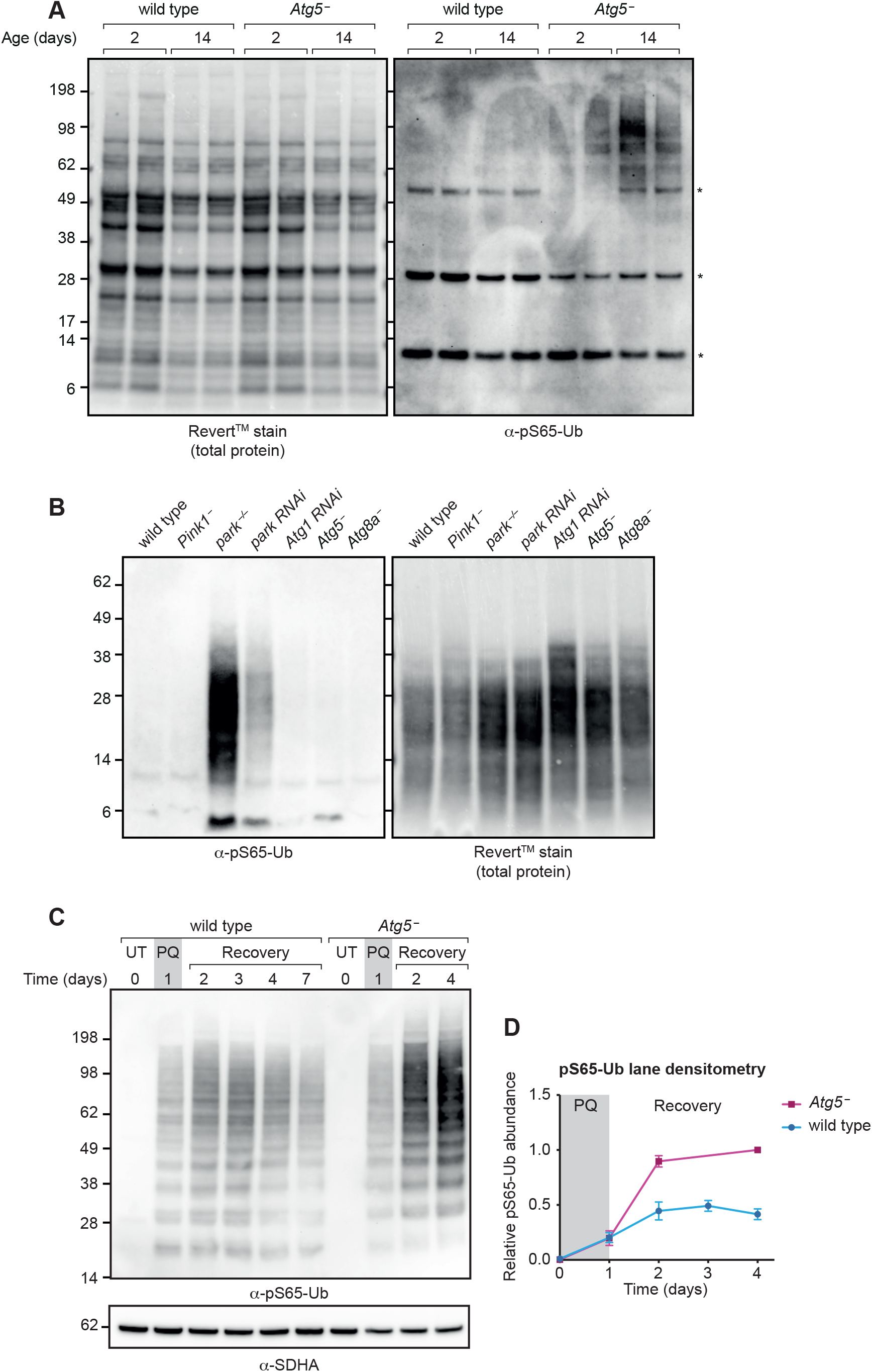
(A) pS65-Ub immunoblot of mitochondrial fractions from wild-type and *Atg5*^−^ flies harvested at the indicated ages. * = non-specific band. (B) pS65-Ub immunoblotting in whole-animal lysates from wandering L3 larvae of the indicated genotypes. (C) pS65-Ub immunoblot of mitochondrial fractions from wild-type and *Atg5*^−^ flies following a paraquat (PQ) pulse-chase assay. UT, untreated; Recovery, return to normal food. (D) Quantification of pS65-Ub lane densitometry from n = 3 independent replicates of (B), expressed relative to the most intense band in each blot. Charts show mean +/− SEM.

